# Long-term, age-associated activity quantification in the DE50-MD dog model of Duchenne muscular dystrophy (DMD)

**DOI:** 10.1101/2025.03.26.645481

**Authors:** Kamila Karimjee, Rachel C.M. River, Emil Olsen, Yu-Mei Chang, Dominic J. Wells, Monica A. Daley, Richard J. Piercy

**Author notes:** IVC Evidensia, Strömsholm Specialist Animal Hospital, 73494 Strömsholm, Sweden.

## Abstract

Animal models with a clinically relevant phenotype remain important for robust evaluation of novel therapeutics for the fatal, X-linked genetic disorder, Duchenne Muscular Dystrophy (DMD). Demonstration of functional improvement is crucial for both patients and regulatory authorities. DMD is associated with a decline in musculoskeletal function with progressive paresis, muscle atrophy and fibrosis: phenotypic features that are also seen in the DE50-MD canine model of DMD. Here we investigate non-invasive methods to quantify changes in activity and behaviour in DE50-MD dogs, using collar-based, tri-axial accelerometers. We measured activity in affected DE50-MD male dogs (3-8 per age point) and littermate wild-type (WT) male controls (3-13 per age point) at monthly intervals from 3 to 18 months of age using Axivity-AX3 accelerometers attached ventrally on each dog’s collar. Data were recorded for 48 hours while dogs remained in their kennels with outside runs following their normal routine. Acceleration vector magnitudes were used to derive various activity indicators over a 24-hour period. Mixed model analyses were used to examine differences between affected and WT groups at different ages. DE50-MD dogs’ activity indicators were significantly higher for % time spent at rest (p<0.001) and significantly lower for all other activity indicators (all p<0.05), when compared to age-matched WT dogs. Sample size calculations reveal that these non-invasive and objective biomarkers offer significant promise for preclinical testing of therapeutics in this model of DMD. Our approach reveals opportunities for cross-model standardisation of activity monitoring methods, applicable to both research and companion animal settings.

**Summary statement:** The DE50-MD dog model of Duchenne muscular dystrophy shows significant age-associated reduction in activity quantified through non-invasive, wearable accelerometers. Activity metrics tested show promise for objective assessment of activity patterns for preclinical trials.

## INTRODUCTION

The X linked disorder, Duchenne Muscular Dystrophy (DMD), affects approximately 1 in 3500-6000 live male births worldwide (Dooley et al., 2010, Mendell et al., 2012, Mercuri and Muntoni, 2013). Certain treatments are licenced or are in clinical trials for DMD, however at present, there is no cure and affected boys typically must use a wheelchair by their early teens (Vuillerot et al., 2010, Bello et al., 2016, Singh et al., 2018); patients die of cardiac and respiratory failure in their twenties or early thirties (Eagle et al., 2002, Nigro, 2012). Consequently, novel therapeutics are urgently needed: these generally require testing in animal models.

There are various animal models of DMD including the mdx mouse (Bulfield et al., 1984) and several canine models (Shimatsu et al., 2005, Walmsley et al., 2010). When compared to the mdx mouse, dog models display a phenotype that better reflects that of affected humans, both functionally and histologically. A widely characterised canine model is the GRMD dog which has a mutation in intron 6 of the dystrophin gene (Valentine et al., 1986, Cooper et al., 1988). The DE50-MD dog is an alternative canine model that provides an additional platform for translational research. These animals have a splice site mutation in the dystrophin gene “hotspot” for human DMD, leading to absence of exon 50 in mature transcripts, a premature stop mutation and absence of dystrophin in striated muscles (Walmsley et al., 2010). Affected dogs have typical dystrophic features including progressive paresis and skeletal muscle atrophy that has been characterised histologically (Hildyard et al., 2022) and by MRI (Hornby et al., 2021). Various inflammatory biomarkers have been examined in this model (Riddell et al., 2021, Riddell et al., 2022) and we previously have used this model to reveal dystrophin restoration following adeno-associated (AAV) virally-mediated systemic CRISPR/Cas9 gene editing (Amoasii et al., 2018).

Functional assessments are crucial for monitoring disease progression in patients with DMD and for monitoring treatment trials: they provide important information on the effects of musculoskeletal changes on overall mobility, as well as assessing quality of life, both of which are critical for patients and their families; they are also important for regulatory bodies. There are several commonly used functional outcome measures in DMD-affected boys including the 6-minute walk test (6MWT) (McDonald et al., 2010), North Star Ambulatory Assessment (Ricotti et al., 2016) and Brooke Upper Extremity Scale (Brooke et al., 1981). Following its success in human assessment, the 6MWT has now become the quantitative, functional outcome measure of choice for assessing disease progression in canine DMD models (Acosta et al., 2016). However, there are several limitations associated with the use of the 6MWT in dogs, just as there are in humans. The original guidelines issued for the test in adult humans included a very rigid protocol outlining specific instructions for limited encouragement and assessor involvement. These guidelines were then adapted as needed for use in children. However, for dogs, there are no validated instructions available for an ‘acceptable’ level of interaction between either the tester or handler and the animal: a dog’s individual response to the test might vary between days and the handler must be adaptive to encourage completion of the test. Further, typically these tests are performed with dogs on leads, meaning that the test result might be influenced by a handler’s walking speed and direction, and the compliance of the animal. In addition, the 6MWT test in dogs is subject to bias, as dog handlers are sometimes unavoidably aware of the genotype of the animal being tested and as such, might handle dogs differently. These factors combined, make interpreting data from the 6MWT challenging. The limitations highlight a need to develop quantitative and objective tools for mobility and activity evaluation in everyday settings that minimise bias and handler constraints in animal models.

Physical activity declines in boys with DMD as their disease progresses (Ricotti et al., 2023). Overnight activity has been assessed in the GRMD dog model using video monitoring: researchers detected reduced activity in affected versus control dogs (Shin et al., 2013, Hakim et al., 2015). We hypothesised that the same is true in the DE50-MD dog model; however, long-term video monitoring typically requires extensive manual labelling of captured video, which might also be subject to observer bias, is time consuming and requires significant digital storage infrastructure.

An increasingly popular method of quantifying movement patterns in humans is via wearable, tri-axial accelerometers (Farrahi et al., 2019). Accelerometer-based methods for assessment of gait have been explored in the GRMD canine models of DMD (Barthelemy et al., 2009).

The use of wearable sensors for tracking changes in longer-term activity in boys with DMD has increased (Gonzalez Barral and Servais, 2025). Recent approval of Stride Velocity 95^th^ centile (SV95C) as a primary endpoint for human clinical trials highlights the shift towards inclusion of and usefulness of digital biomarkers in assessing functional outcomes (Servais et al., 2024). However, unlike in boys, tracking changes in longer-term activity has not yet been explored in canine DMD models. Moreover, consensus has not yet been reached as to the most appropriate aggregate metrics to report activity data for dogs: different groups report varied methods, and without derivation instructions (Hansen et al., 2007, Yam et al., 2011, Clarke and Fraser, 2015, Griffies et al., 2018). This makes it very difficult to compare outputs between studies and to justify the choice of metrics, or to replicate work. This issue was discussed in detail in previous work (Karimjee et al., 2024) in which we explored a standardised, open-source approach to canine activity monitoring to make studies comparable and to determine robust outcome measures for quantifying disease progression and, potentially, to evaluate novel treatments. In this current longitudinal study, we describe use of collar-based accelerometers and this open-source approach to quantify changes in activity patterns with disease progression to characterise the DE50-MD phenotype and to identify suitable biomarkers for planned pre-clinical trials.

## METHODS

### ANIMAL HUSBANDRY

Carrier DE50-MD female dogs were mated naturally with wild type (WT) beagle males (RCC strain). Adult dogs were housed (12-hour light/dark cycle; 15-24°C) in large kennels in groups, until pregnant females were close to whelping when they were housed in single kennels. Puppies were weaned at 10-12-weeks-old, then grouped in their litters until approximately 4 to 5 months of age. Adults were then housed in groups of 2-4 animals according to social hierarchy. Kennel size ranged between 6.2 – 6.6 m^2^. Dogs were fed Burns (Burns Pet Nutrition Ltd, Wales, U.K.) puppy or adult feed, as required, twice a day ad lib, with daily human interaction and access to outdoor runs and grassy paddocks (approximately 100m^2^) in groups of up to 5 dogs, typically between the hours of 8am – 3pm. Husbandry conditions exceeded the minimum Animal (Scientific Procedures) Act (1986) (ASPA) requirements. Puppy genotypes were confirmed by sequencing of PCR products amplified from cheek swab-derived DNA (Walmsley et al., 2010) within 7 days from birth and corroborated by measurement of serum creatine kinase activity (data not shown) (Riddell et al., 2021). Unaffected animals, not used for research or breeding, were rehomed at around 12 weeks of age or at study termination. Carrier females underwent routine ovariohysterectomy prior to rehoming.

### ARRIVE GUIDELINES

ARRIVE guidelines were followed for the design and conduct of the study. Animals were assigned to groups based on their genotype (WT vs DE50-MD) with the sample sizes as summarised in Figure 1 with inclusion based on availability over the experimental period. There were no specific exclusion criteria and randomisation was not conducted as there was no treatment group and hence no outcome measurements. Specific blinding was not performed as all data collection was obtained objectively through use of accelerometers. Statistical methods are summarised in the data collection section below. All experimental procedures involving animals in this study were conducted according to UK legislation within an ASPA project licence (P9A1D1D6E), however activity monitoring work was sub-threshold. All efforts were made to minimise any animal suffering throughout the study. Pre-determined end-points for DE50-MD dogs were established including dehydration (unresolved by fluid treatment), lethargy/motor dysfunction, weight loss/dysphagia, dyspnoea, listless behaviour/demeanour, or heart failure. All dogs were observed at least twice daily by animal technicians; animal displaying any of these signs were reported to and assessed by the Study Director, the Named Veterinary Surgeon (NVS) and the Named Animal Care and Welfare Officer (NACWO). Should any dog reach any of the pre-determined end-points, prior to the planned 18-month study end, they were humanely euthanised. Euthanasia was performed using an overdose of sodium pentobarbital (250 mg/kg, Dolethal, Covetrus) administered intravenously via preplaced catheter. Of the 12 DE50-MD dogs included in this study, 4 DE50-MD dogs (DE50-AH3, -T4, -T6, -V2) were euthanised prior to the end of the study due to reaching pre-determined humane end-points (all related to dysphagia – no other humane endpoint was reached during this study). All 18 WT dogs were rehomed.

**Figure 1:**
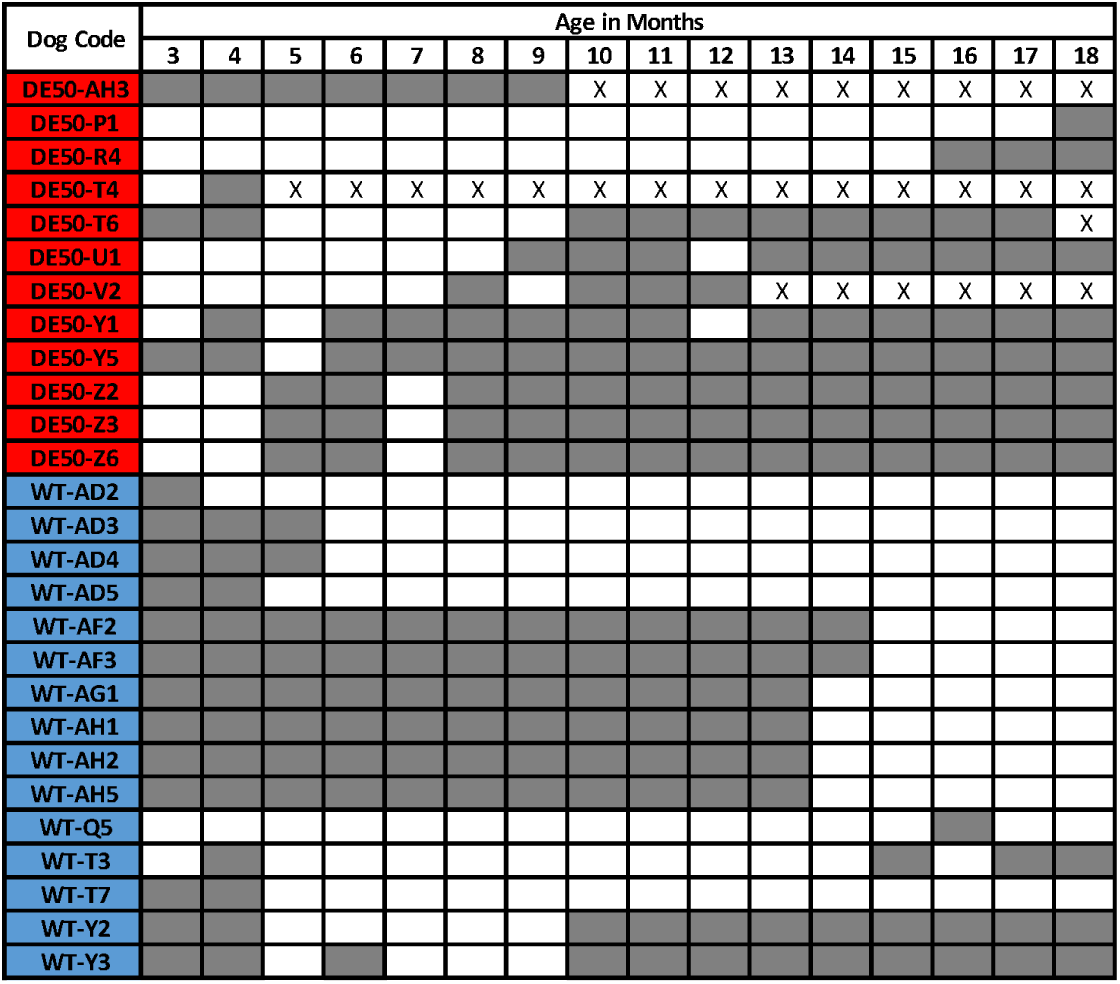
Summary of dogs included in this study and ages sampled. Each row corresponds to an individual dog and each column corresponds to the dogs’ ages in months. The dog code is derived from the genotype: DE50 (red) or WT (blue), the following letters refer to the litter and the final number refers to the order of birth of the puppy, e.g. the first puppy born would be labelled 1, the second labelled 2 etc. Grey shading indicates that the dog was sampled at the indicated age. A white square indicates that data were not collected from that dog at that age. An X within a white square indicates that data were not collected from that dog due to euthanasia prior to the time point.

### DATA COLLECTION

Data were collected via Axivity AX3 activity monitors (Axivity, U.K.) which have previously been validated for use in dogs (Kiyohara et al., 2015, Ladha et al., 2017, Griffies et al., 2018, Karimjee et al., 2024) as their parameters are easily modifiable by the user through open-source software. The devices were mounted onto each dog’s neck collar using duct tape, with the logger positioned ventrally, to minimise the likelihood of the collar’s rotation. However, note that as the selected metrics are direction invariant, rotation of the device was not relevant. Dogs were habituated to wearing collars from 7 weeks of age. Dogs were sampled each month between the ages of 3 and 18 months old. Practical constraints meant that not all dogs were sampled at every age point (summarised in figure 1): this work was part of a wider study and selection criteria for inclusion in this study was based on availability of animals; this led to variation in the number of individuals sampled at each time point. All DE50-MD and WT dogs were male.

Weekday monitoring was for 48 hours continuously (weekdays 12pm-12pm) at a sample frequency of 200-400 Hz starting at 12pm. Where sample frequencies greater than 200 Hz were used, data were computationally resampled to 200 Hz to ensure homogeneity. Dogs remained in their kennels (with their kennel mates) during the test period and were subject to their “normal” daily routine (feeding, cleaning etc). To best characterise each time point, the 48-hour time interval, beginning at 12pm on the first day of monitoring, was split into 2 x 24-hour periods enabling aggregate metrics to be calculated per 24 hours. The mean of these metrics across the two, 24-hour periods was calculated and used in subsequent analyses. Variability between the two 24-hour periods was assessed and is described in further detail below. In some cases (15/214 recordings), it was not possible to capture 48 hours of continuous activity due to sensor destruction (chewing) or malfunction. In these cases, aggregate metrics were calculated for the first 24-hours.

### SIGNAL PROCESSING

All signal processing was performed in MATLAB R2020b. A 6^th^ order Butterworth band-pass filter was used in both directions to obtain zero phase lag. Cut-off frequencies of 0.28 Hz and 32.76 Hz were used, based on pilot observations (Karimjee et al., 2019). The vector magnitude was computed from the filtered data using the formula below.

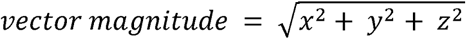

The vector magnitude was smoothed using a 0.3 second epoch length and activity aggregate metrics were then calculated (see (Karimjee et al., 2024) for full derivation) from the vector magnitude as described below.

#### DAILY ACTIVITY INTENSITY

This metric characterises overall, whole body acceleration over 24 hours and is calculated by computing the integral of the 24-hour vector magnitude curve minus gravity. As this calculation takes place after band-pass filtering, the signal due to gravity has already been removed by the high pass component of the filter. Daily activity intensity is a velocity quantity (in meters per second: ms^-1^).

#### MX_ACC_

These metrics are defined as the acceleration threshold that defines the animal’s X most active minutes, accumulated over a 24-hour period. M2_ACC,_ M30_ACC,_ M60_ACC_ values were characterised according to work outlined in (Karimjee et al., 2024). Briefly, advantages of these metrics include producing data that are comparable between study populations without adjustment or correction. They can also be complimented with relevant, species-specific activity intensity levels (e.g. walking, trotting) from reference data, thereby facilitating interpretation and providing additional real-world context (Rowlands et al., 2019). Note, these reference thresholds correspond to acceleration intensity levels exhibited during different gaits by healthy dogs in previous work (Karimjee et al., 2024): they were not used to detect specific gait types in this study.

#### PERCENT TIME SPENT INACTIVE

This metric quantifies the proportion of time spent at rest over a fixed period derived from our previously labelled accelerometer and video data and Receiver Operating Characteristic (ROC) analyses (Metz, 1978). We utilised a threshold (that maximises sensitivity and specificity) of 0.154g and the proportion of time spent below that threshold was calculated.

#### PERCENT TIME SPENT AT “HIGH” INTENSITY ACTIVITY

This metric quantifies the proportion of time spent at a higher intensity activity level, defined as the threshold that best discriminates between DE50-MD and WT control dogs. This threshold was derived by computing the percentage of time spent above an acceleration threshold, in a similar way as described in (Karimjee et al., 2024) to compute the percent time spent active. However, to determine the most appropriate threshold that best discriminated between groups, we repeated this computation, iterated through threshold values between 0.05g and 1.0g in increments of 0.01g and calculated the percent time spent above each chosen threshold per recording. This resulted in 100 complete data iterations, with a datapoint per animal (% time spent above each threshold) per age point per threshold used. ROC analysis was then used for each threshold: we computed the area under the curve (AUC) and chose the first threshold (0.755g) after which the AUC value stopped increasing (0.999), and corresponding to the dataset computed with the threshold parameter that best discriminated DE50-MD and WT dogs.

#### PERCENT TIME SPENT AT LOWER INTENSITY ACTIVITY

This metric quantifies the proportion of time spent at a lower intensity activity, which was defined as being between the two thresholds described above (above 0.154g and below 0.755g) activity over the fixed period.

### STATISTICAL ANALYSES

Statistical analyses were conducted with SPSS Statistics (IBM SPSS Statistics 25) and GraphPad Prism 9 (GraphPad Software Inc. 1994-2021). To assess the validity of averaging the activity metrics across the two 24-hour periods of data, we compared Daily Activity Intensity data from the 1^st^ 24-hours with the 2^nd^ 24-hour for all dogs at all time points. Comparison across the 1^st^ and 2^nd^ 24-hour periods was performed using a two tailed, paired t-test. The correlation between Daily Activity Intensity in the 1^st^ and 2^nd^ 24-hour periods was assessed by linear regression.

A principal component analysis (PCA) was performed (GraphPad Prism 9) to summarise variation in activity metrics among the DE50-MD and WT controls across all ages. This allowed us to explore which activity metrics varied most across all dogs and assess whether these patterns differed between groups. The first component (PC1), which captured the largest proportion of variance, was then examined statistically using a linear mixed-effects model (LMM), assessing the effect of age, genotype and their interaction, followed by Fisher’s Least significant difference post-hoc comparisons. Individual dog was included as a random effect to account for repeated measures. Individual activity metrics were assessed using the same LMM analysis as described above (SPSS Statistics).

Sample sizes were calculated to assess the metrics that would be most appropriate for assessing treatment outcomes in possible future pre-clinical treatment trials (power 0.8, alpha 0.05). Effect sizes of 20%, 25%, 50%, 75% and 100% were investigated (i.e. the degree to which any therapy could induce a change from the DE50-MD value towards that of WT animals). Sample size calculations were completed using GLIMMPSE v1 (Kreidler et al., 2013) taking into account repeated measurements within individuals. Since GLIMMPSE only allows for 10 repeated measures, sample size calculations were completed using data from 3 month intervals, rather than monthly. Ages included were 3, 6, 9, 12, 15 and 18 months which correspond with the same intervals used in other phenotypic analyses by our group in dogs from the same colony (Hornby et al., 2021, Riddell et al., 2021, Hildyard et al., 2022, Riddell et al., 2022).

## RESULTS

There was no significant difference in Activity Intensity between the 1^st^ and 2^nd^ 24-hour periods across all ages for WT (p>0.1; Figure S1A) or DE50-MD (p>0.1; figure S1B) dogs; further, a significant correlation between the two 24-hour periods existed for each genotype (p<0.0001; WT: slope estimate: 0.75 +/- 0.09 SE; DE50-MD slope estimate: 0.69 +/- 0.08 SE; supplementary figure S1C). Consequently, further analyses were typically performed using the mean of the 24-hour periods for all activity metrics for each animal.

Activity metrics are summarised in figure 2 and supplementary table 1. Percent time active was greater across all ages in WT controls compared to DE50-MD dogs and reduced in both groups with age. This effect was seen in both the measured percent time active in low intensity (Fig. 2b) activity (p<0.001 for both age and group) and high intensity (Fig. 2c) activity (p<0.01 for age and group). The interactions between group and age were also significant, such that, in general, the differences between the two groups increased with age. For example, DE50-MD dogs spent 0.68% of their time in high intensity activity at 3 months, reducing to 0.16% at 18 months, compared to WT controls with 1.95% at 3 months and 1.6% at 18 months. The same effects were also seen in the activity intensity (Fig. 2a) metric (all p<0.001).

**Fig. 2:**
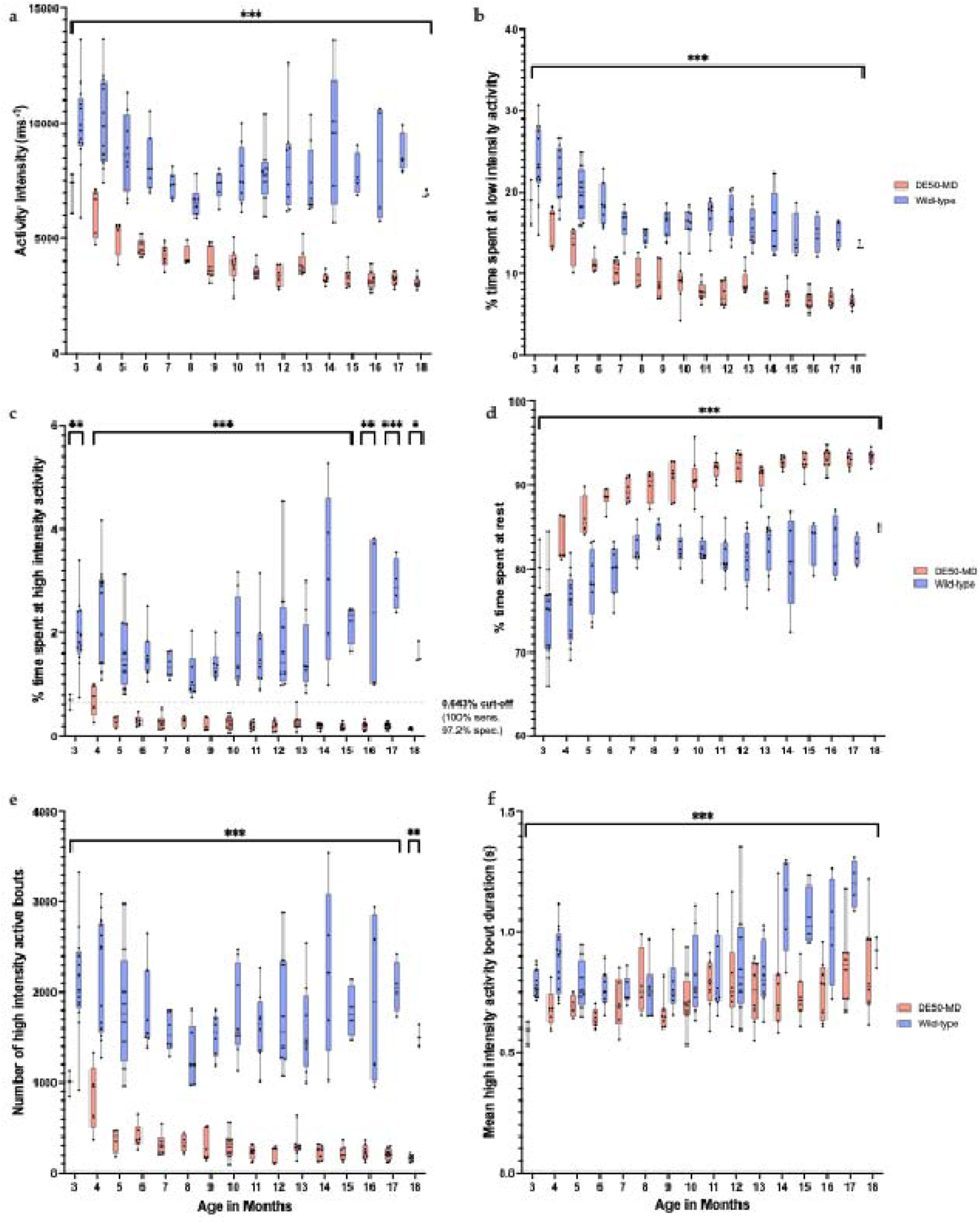
Selected activity metrics for all dogs ages 3-18 months old. From top left to bottom right: a – activity intensity, b – percent time spent at low intensity activity, c – percent time spent at high intensity activity, d – percent time spent at rest, e – number of high intensity active bouts, f – mean high intensity active bout duration. *p < 0.05, ** p < 0.01, *** p < 0.001.

While the number of low intensity active bouts decreased in both groups with age, the mean duration of low intensity active bouts increased in WT controls with age but decreased in DE50-MD dogs (interaction p<0.001). The variation between individual dogs within the DE50-MD group also reduces with age in this metric. The number of high intensity bouts (Fig. 2e) also decreased with age in DE50-MD dogs, but remained relatively consistent in WT dogs, with a significant difference between groups at all ages (p<0.001).

The percent time spent resting (Fig. 2d) was greater across all ages in DE50-MD dogs compared to WT controls and increased in both groups with age (age and group: p<0.001). The interaction effect between age and group was also significant, the difference between groups in this metric increased with age (p<0.001).

The threshold above which the dogs’ X most active minutes were accumulated over a 24-hour period (X values of 2, 30 and 60 mins) were greater across all ages in WT controls compared to DE50-MD dogs (p<0.01) (figure 3; supplementary table 2). We observed that for all dogs (DE50-MD and WT), the 2 most active minutes over the 24-hour period (M2_ACC_) are at an intensity level greater than standing and walking. For the WT dogs, M2_ACC_ values were also at an intensity level much greater than trotting. In contrast, DE50-MD dogs’ M2_ACC_ values were closer to the intensity level of trotting, with one individual below the trotting reference value. The WT dogs’ M30_ACC_ values were all greater than the walk reference value; hence all WT dogs carried out 30 minutes of activity of intensity greater than or equal to walking, with many individuals surpassing the trotting threshold. However, for the DE50-MD dogs, the M30_ACC_ values were almost all below the trotting intensity threshold (except one individual), with only some individuals reaching 30 minutes of activity equivalent to intensity of walk. When examining the dogs’ M60_ACC_ values, some of the DE50-MD dogs’ 60 most active minutes fall below the walk intensity level. This contrasts with WT dogs, where all dogs carry out 60 minutes of activity greater than or equal to walking intensity, with many surpassing the trotting intensity threshold. There was an increase in MX_ACC_ values with age in WT controls, and a decrease in DE50-MD dogs, however this was only significant for X values of 30 and 60 mins (p<0.01). Generally, the differences between groups increased with age in all metrics, supported by a significant interaction effect between age and group (all p<0.01).

**Fig. 3:**
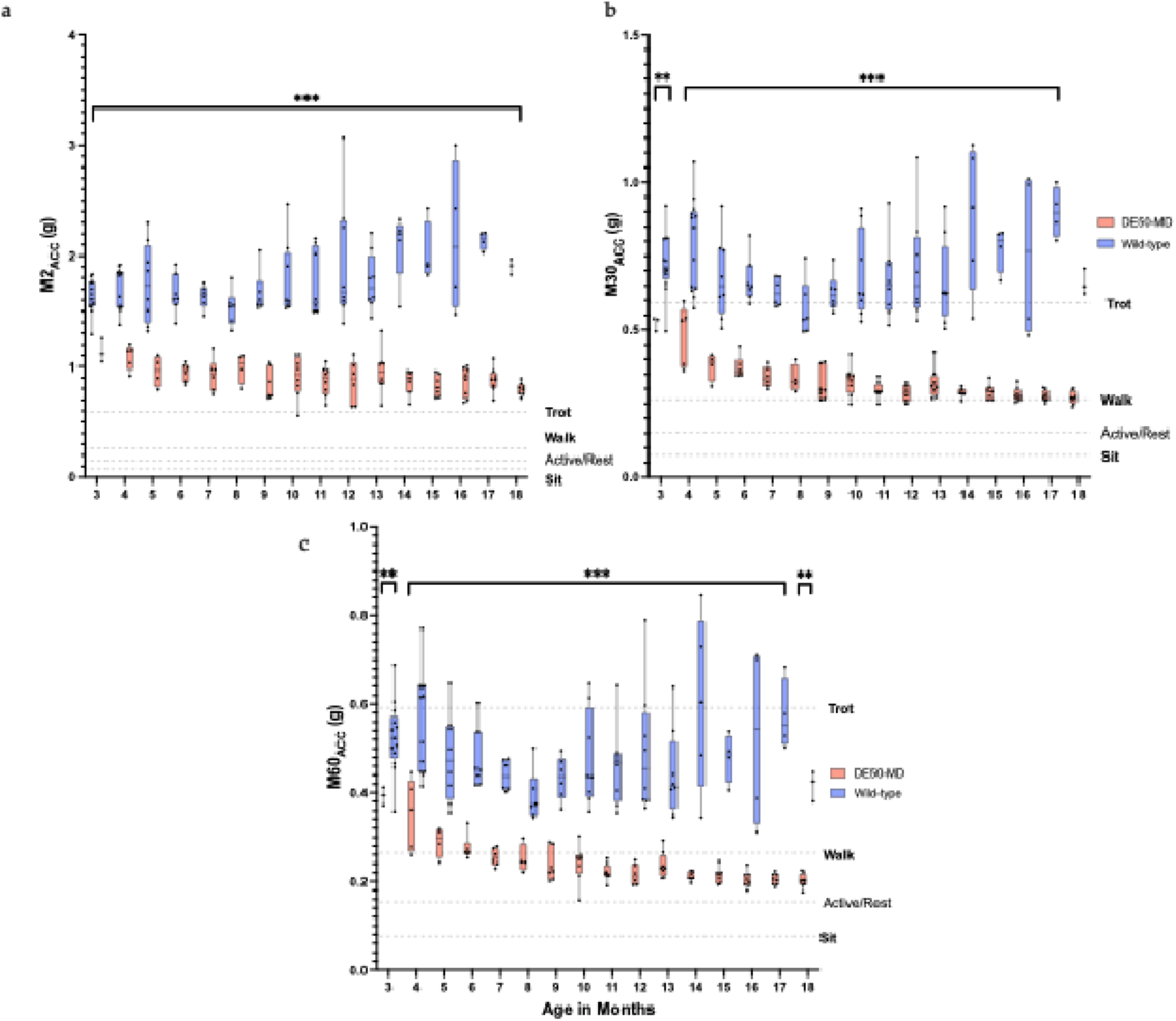
a - MX_ACC_ activity metrics for all dogs ages 3-18 months. M2_ACC,_ b - M30_ACC_, c - M60_ACC_ activity metrics for all dogs between 3 and 18 months old. MX_ACC_ metrics quantify the threshold above which the dog’s X most active minutes are accumulated over a 24-hour period. Reference activity levels for locomotor behaviours of interest included on each graph for context. *p < 0.05, ** p < 0.01, *** p < 0.001.

**Fig. 4:**
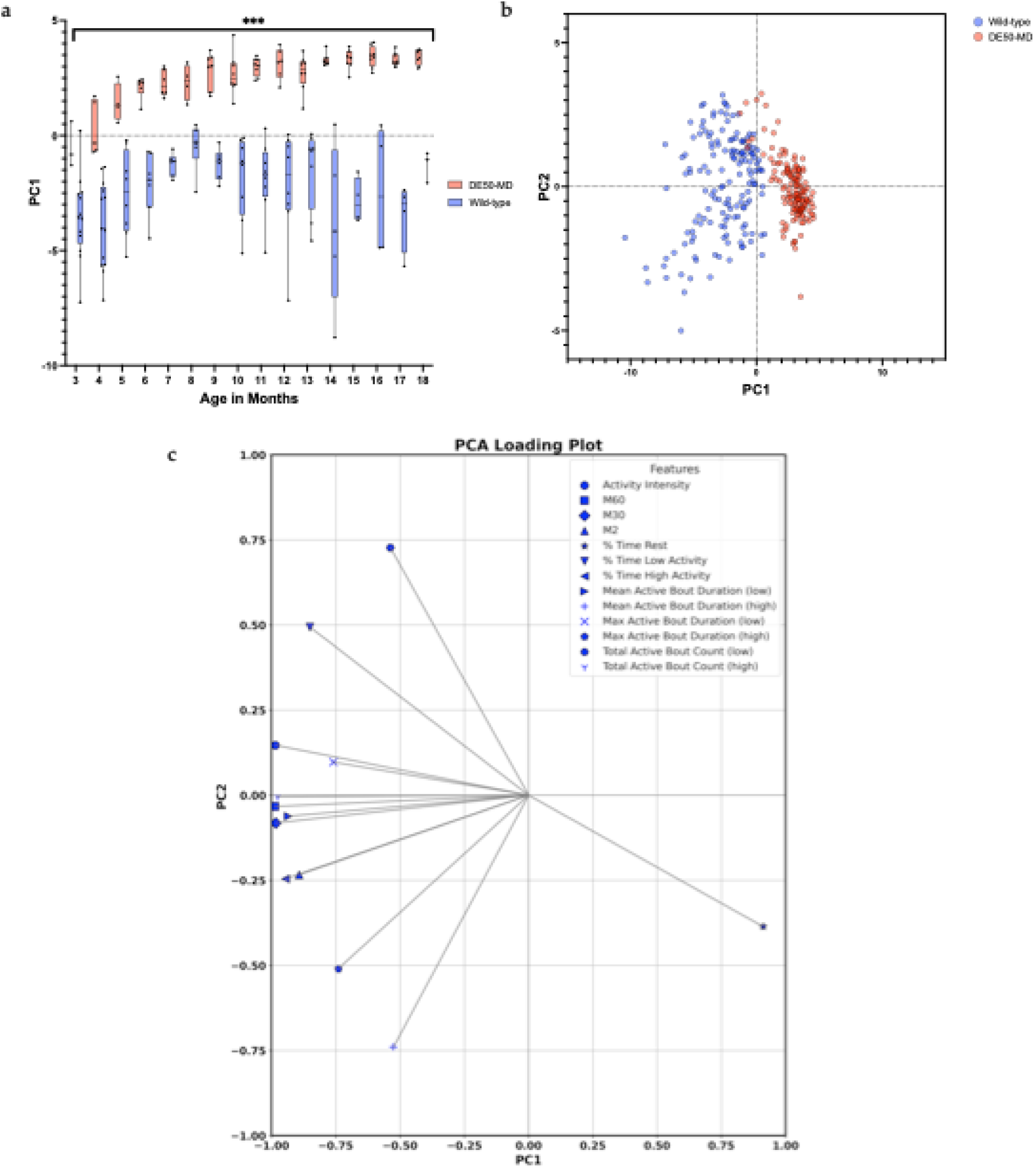
PCA analysis outputs a) PC1 by age in months for both groups, b) biplot, c) loadings. a – Principal component 1 data for all dogs between 3 and 18 months, *p < 0.05, ** p < 0.01, *** p < 0.001, b – biplot showing dogs in PC1 and PC2 space, c – loading plot showing contribution of variables to PC1 and PC2.

PCA revealed two principal components (PCs) of interest with Eigenvalues of 10.53 and 2.02 which cumulatively explained 89.67% of the total variance (75.21% and 14.46% respectively). When examining the loadings of the two PCs, time spent resting, and metrics quantifying higher intensity activity contribute most towards PC1 and the number of low intensity activity bouts contributed most to PC2. Therefore, positive PC1 scores correspond to higher percent time spent at rest and negative PC1 correspond to higher levels of activity. Positive PC2 scores then indicate more time in low intensity activity, and negative PC2 scores indicate more time at higher intensity activity. PC1 scores were greater in DE50-MD dogs at all ages when compared to WT controls (p<0.001) and generally increased in both groups with age (p<0.001) with the differences between groups also increasing with age (interaction effect p<0.001). There were no significant differences between groups in PC2 scores, however there was a decrease in scores with age (p<0.001).

We aimed to determine which of the metrics examined in this work would be most applicable to assessment of activity patterns in pre-clinical trials. We calculated prospective required sample sizes (Table 1) for all activity metrics that showed statistically significant differences (p < 0.05) when discriminating between genotype groups. Calculations were computed for all metrics at 3, 6, 9, 12, 15 and 18 months old taking into account repeated measures within individuals. The number of study animals required to detect an improvement of 100%, 75%, 50%, 25% and 20% towards WT levels was quantified. Result show that all metrics show promise to discriminate between groups; indeed, of all metrics computed, the total number of bouts at high intensity, M2 _ACC,_ M30_ACC_ and PC1 metrics can detect as little as 20% effect with low numbers of animals (up to 6 per group).

**Table 1:** Sample size calculations for all statistically significant metrics at selected ages. Calculations (power 0.8, alpha 0.05) completed for all activity metrics that showed statistically significant differences between groups by effect size. Numbers represent the number of animals required per genotype to detect the desired treatment effect at this statistical power in future trials using each metric. Note that the total bout count of high intensity activity and M30_ACC_ metrics can detect as little as a 20% treatment effect with very few animals (five). The combined approach (via PCA) tends to result in fewer animals needed to detect any given difference.

| Activity Metric | Group size to detect change (N per genotype) |  |  |  |  |
| --- | --- | --- | --- | --- | --- |
|  | 100% | 75% | 50% | 25% | 20% |
| Activity Intensity<br>(ms<sup>-1</sup>) | 2 | 2 | 3 | 5 | 6 |
| % time spent at rest | 2 | 2 | 3 | 6 | 8 |
| % time spent at low intensity | 2 | 3 | 3 | 7 | 9 |
| % time high intensity | 2 | 2 | 3 | 5 | 6 |
| Avg. bout duration low intensity (s) | 3 | 3 | 5 | 13 | 19 |
| Total bout count low intensity | 3 | 4 | 6 | 17 | 26 |
| Avg. bout duration high intensity<br>(s) | 4 | 5 | 9 | 33 | 51 |
| Total bout count high intensity | 2 | 2 | 3 | 4 | 5 |
| M2<sub>ACC</sub> (g) | 2 | 2 | 3 | 4 | 6 |
| M30<sub>ACC</sub> (g) | 2 | 2 | 2 | 4 | 5 |
| M60<sub>ACC</sub> (g) | 2 | 2 | 3 | 5 | 7 |
| PC1 | 2 | 2 | 3 | 4 | 6 |

## DISCUSSION

To our knowledge, this is the first study that has investigated long-term changes in activity patterns with accelerometry in a dog model of DMD. Previous functional measures have included gait kinematics (Barthélémy et al., 2011, Shin et al., 2013), walking distance assessments (Acosta et al., 2016, Pizzato et al., 2016) and video monitoring (Shin et al., 2013). Although these studies quantify functional performance in various ways, they all report globally-reduced mobility in affected dogs when compared to controls. The primary aim of our work was objectively and quantitatively to measure the activity phenotype of the DE50-MD canine model, using non-invasive methods that minimise bias. DE50-MD and age-matched, littermate WT controls were tested monthly between 3 and 18 months of age and we quantified intensity of activity using both acceleration- and time-based thresholding techniques, each of which has advantages (Karimjee et al., 2024). Briefly, the combination of these analysis methods allows for interpretation of different aspects of activity patterns alongside each other, including subtle differences in function that would not be seen if any of the metrics was used alone.

The ability to distinguish between intensity levels of activity when quantifying treatment effects would likely provide insight into how improvements in function might affect quality of life. For example, an increase in percent time spent at lower intensity activity only and no change in higher intensity activity could be a valuable outcome, as well as providing a more granular understanding of how drug mechanism of action is reflected in functional outcomes. Our overall goal was to determine the most useful quantitative metrics to distinguish between the activity patterns of the DE50-MD and WT controls for future preclinical trials of therapeutics. These data reveal the prominent activity differences between DE50-MD dogs and WT dogs across all ages studied. Time spent resting and metrics quantifying higher intensity activity appear to be key factors. Also importantly, these metrics are likely to be helpful as objective functional outcome measures, using low animal numbers, in future treatment trials (see Table 1) for detection of even low to moderate phenotypic improvements, which if translated to humans, could have massive impacts on quality of life.

Measures of activity intensity (ms^-1^) provide global functional indicators of physical activity by quantifying the total change in velocity over 24-hours. We observed a slow decline in this metric with age, in line with disease progression (Hornby et al., 2021). A slight decrease in activity intensity was also seen in the WT controls with age, but unlike the affected dogs, this plateaus at 8-9 months. There was lower intra-class variation in the DE50-MD group across ages when compared to the WT controls. The increased variation in WT control groups was greater at older ages, suggesting a larger range of activity levels but this might be due to lower sample size of animals at these older ages. These findings are in line with previous work quantifying the gait phenotype of the GRMD dog model. There, gait parameters including stride frequency, and speed of voluntary movement were reduced in GRMD dogs compared with controls (Marsh et al., 2010, Barthélémy et al., 2011) although those studies were trial-based protocols, as opposed to continuous collection over a longer periods as done here. Nonetheless, the observed differences in the gait parameters seen in the GRMD dogs support the findings of reduced levels of spontaneous activity in dystrophic dogs.

The proportion of time spent at rest quantifies time spent under the pre-determined “active/inactive” threshold for acceleration (<0.154g) over a 24-hour period (Karimjee et al., 2024). As expected, this was greater in DE50-MD dogs at all ages when compared to WT controls. The proportion of time spent resting increased steadily with age in the DE50-MD group, but plateaued at 8-9 months in the WT control group, the latter potentially coinciding with young adulthood as Beagles reach 95% of their final body weight at 246 days (8 months) (Salomon et al., 1999).

Reductions were observed in all metrics across both high and low intensities, however the most prominent differences are seen at higher intensity levels, particularly the proportion of time spent at high intensity activity and the total number of high intensity bouts of activity over a 24-hour period. These metrics decreased over time, particularly in the DE50-MD group. This followed the pattern of expected disease progression in human patients, where individuals undergo progressive paresis, with stiffer muscles (Lacourpaille et al., 2015) and reduced range of joint motion (de Vocht et al., 2019). These same factors might influence voluntary spontaneous high intensity movements in DE50-MD dogs, however relative paresis might also play a role. This was also supported by other group members’ work that recently reported reduced muscle volume (Hornby et al., 2021), histopathological changes (Hildyard et al., 2022) and enhanced eccentric contraction-induced force decrement of tibiotarsal flexor muscles (Riddell et al., 2018, Riddell et al., 2023) in DE50-MD dogs when compared to WT controls. Similar results were also seen in the GRMD dog model (Childers et al., 2002).

In addition to more traditional acceleration threshold metrics that are derived to be specific to a certain experimental population, we have assessed time-based threshold metrics: M2_ACC,_ M30_ACC_ and M60_ACC._ These represent the acceleration levels (measured in units of g) above which the dog’s X most active minutes are accumulated over a 24-hour period. Use of these metrics can facilitate direct comparisons among and between disease models and even species. This is an important benefit when considering data from different pre-clinical models of DMD. As well as observing differences between groups, these metrics can be used to compare the intensity levels of activity to reference values, derived from earlier work (Karimjee et al., 2024) collected from a cohort of 6 healthy animals contained within this dataset. Overall, whilst the MX_ACC_ indices provide insight into the intensity of activity relative to relevant canine-specific movements, it is important to note that these results do not indicate a tendency or preference for a particular gait type. The comparisons are between the measured intensity levels and known reference behaviours in healthy animals. It is possible, and even likely, that the accelerations measured for specific behaviours such as walking could differ between DE50-MD and WT dogs. Future work, aimed at better classifying specific activities that contribute to these metrics, will enable a richer assessment of behavioural differences between groups. However, these metrics are valuable in enabling quantitative comparisons of activity intensities between different canine groups and disease models, providing a consistent and objective assessment tool to use across different research groups.

Sample size assessments were performed to help determine the most useful activity biomarkers to take forward to pre-clinical trials in the DE50-MD dog model. The number of high intensity active bouts over a 24-hour period and the M30_ACC_ (the threshold above which the dog’s 30 most active minutes are accumulated over 24-hours) had the lowest number of animals required for the smallest treatment effect size (20%) across all ages (n=5). Examining the outputs of the PCA also reveals that the combination of our proposed activity metrics can successfully distinguish between the DE50-MD and WT controls. When examining the outputs of the PCA, the loadings of the first two principal component - time spent resting and metrics quantifying higher intensity activity - contributed most towards PC1 and the number of low intensity activity bouts contributed most to PC2. The PC1 statistic also offers promise as a biomarker for assessing responses to treatments in this model.

This study was carried out whilst the dogs remained in their habituated housing, undisturbed. This provides a longer-term assessment of the animals’ capacity for movement, in contrast to related studies involving shorter duration (Barthelemy et al., 2009) which might depend on the animals’ performance at that particular time of day, or influence of the handler. Similarly, methods involving scoring based on interpretation of video footage or direct observation might also be subject to observer bias; our approach minimizes these potential biases.

In summary, quantitative, objective assessment into the activity patterns of the DE50-MD dogs and WT controls have been provided, demonstrating the ability to distinguish between these groups across a range of ages. We have reported a battery of activity metrics derived from tri-axial accelerometer data which can contribute towards characterising the functional performance of the DE50-MD model for DMD as well as discriminating between genotypes. These results support the hypothesis that these activity metrics, collected via long-term activity monitoring, can successfully discriminate between genotypes, and will be useful in assessing treatment outcomes in future pre-clinical trials. The fact that these assessments are easily obtained, are non-invasive and independent of bias has distinct practical, welfare and scientific advantages. Similar approaches might be applicable in other canine models of DMD and indeed, more generally, in other animal disease models associated with locomotor or neuromuscular dysfunction.

## Acknowledgements

The authors thank colleagues at the Royal Veterinary College for their contributions to this work and staff within the Biological Services Unit at the Royal Veterinary College for excellent and compassionate care of the animals.

