## Supplementary data for "Long-term, age-associated activity quantification in the DE50-MD dog model of Duchenne muscular dystrophy (DMD)"

| Age in Months | % time spent at rest*** | | % time spent at low intensity*** | | % time high intensity** | | Avg. bout duration low intensity (s)* | | Total bout count low intensity* | | Avg. bout duration high intensity (s)*** | | Total bout count high intensity*** | |
|---|---|---|---|---|---|---|---|---|---|---|---|---|---|---|
| | *DE50* | *WT* | *DE50* | *WT* | *DE50* | *WT* | *DE50* | *WT* | *DE50* | *WT* | *DE50* | *WT* | *DE50* | *WT* |
| **3** | 80.5 (2.9) | 74.5 (4.5) | 18.9 (2.8) | 23.6 (4.5) | 0.7 (0.2) | 2.0 (0.6) | 2.1 (0.4) | 2.5 (0.6) | 8175$^{f}$ (1802) | 9005$^{f}$ (1306) | 0.6 (0.5) | 0.8 (0.1) | 995 (147) | 2118 (580) |
| **4** | 83.4 (2.7) | 75.5 (4.0) | 15.9 (2.5) | 22.2 (3.4) | 0.7 (0.3) | 2.3 (1.0) | 2.1 (0.5) | 2.7 (0.7) | 6977 (1649) | 8081 (1471) | 0.7 (0.1) | 0.9 (0.1) | 856 (369) | 2196 (646) |
| **5** | 86.2 (2.5) | 78.4 (3.2) | 13.5 (2.5) | 19.9 (3.2) | 0.3 (0.1) | 1.7 (0.8) | 1.8 (0.4) | 2.2 (0.5) | 6774 (1369) | 8722 (972) | 0.7 (0.0) | 0.8 (0.1) | 366 (134) | 1821 (671) |
| **6** | 88.5 (1.2) | 79.9 (3.1) | 11.2 (1.1) | 18.5 (2.6) | 0.3 (0.1) | 1.6 (0.5) | 1.6 (0.2) | 2.2 (0.5) | 6157 (543) | 7894 (780) | 0.6 (0.0) | 0.8 (0.1) | 407 (136) | 1787 (472) |
| **7** | 89.4 (1.5) | 82.4 (2.0) | 10.4 (1.4) | 16.3 (2.1) | 0.3 (0.1) | 1.4 (0.2) | 1.6 (0.2) | 2.1 (0.2) | 5757 (736) | 7220 (966) | 0.7 (0.1) | 0.8 (0.1) | 319 (124) | 1558 (214) |
| **8** | 89.9 (2.0) | 84.1 (1.3) | 9.8 (1.9) | 14.7 (0.9) | 0.3 (0.1) | 1.2 (0.5) | 1.5 (0.3) | 2.0 (0.1) | 5710 (480) | 7015 (705) | 0.8 (0.1) | 0.8 (0.1) | 330 (95) | 1285 (332) |
| **9** | 90.7 (2.1) | 82.4 (1.7) | 9.1 (2.0) | 16.3 (1.7) | 0.2 (0.1) | 1.4 (0.3) | 1.5 (0.2) | 2.2 (0.3) | 5477 (474) | 7146 (556) | 0.7 (0.1) | 0.8 (0.1) | 278 (164) | 1524 (236) |
| **10** | 90.8 (2.5) | 82.1 (2.2) | 8.9 (2.4) | 16.2 (1.8) | 0.3 (0.1) | 1.7 (0.9) | 1.6 (0.2) | 2.3 (0.4) | 5088 (1227) | 6827 (975) | 0.7 (0.1) | 0.8 (0.2) | 307 (146) | 1763 (497) |
| **11** | 92.0 (1.2) | 81.3 (2.6) | 7.8 (1.1) | 17.0 (2.2) | 0.2 (0.1) | 1.6 (0.7) | 1.4 (0.1) | 2.3 (0.4) | 4933 (715) | 7215 (1013) | 0.8 (0.1) | 0.8 (0.2) | 229 (75) | 1618 (389) |
| **12** | 92.4 (1.7) | 81.1 (3.4) | 7.4 (1.6) | 17.0 (2.6) | 0.2 (0.1) | 1.9 (1.2) | 1.4 (0.1) | 2.4 (0.6) | 4599 (669) | 6885 (922) | 0.8 (0.1) | 0.9 (0.2) | 215 (87) | 1785 (657) |
| **13** | 90.9 (1.7) | 82.8 (3.0) | 8.8 (1.5) | 15.6 (2.4) | 0.3 (0.1) | 1.6 (0.7) | 1.4 (0.1) | 2.4 (0.5) | 5485$^{f}$ (689) | 6323$^{f}$ (1028) | 0.8 (0.1) | 0.8 (0.1) | 305 (144) | 1590 (509) |
| **14** | 92.7 (0.7) | 80.8 (5.5) | 7.1 (0.7) | 16.2 (4.0) | 0.2 (0.1) | 3.0 (1.7) | 1.4 (0.1) | 2.7 (0.8) | 4619 (453) | 6173 (904) | 0.8 (0.2) | 1.1 (0.2) | 223 (75) | 2220 (955) |
| **15** | 92.6 (1.3) | 83.3 (2.8) | 7.3 (1.2) | 14.5 (1.2) | 0.2 (0.1) | 2.2 (0.4) | 1.4 (0.1) | 2.4 (0.3) | 4707 (832) | 6049 (760) | 0.7 (0.1) | 1.1 (0.1) | 225 (84) | 1784 (287) |
| **16** | 93.2 (1.2) | 82.8 (3.8) | 6.7 (1.2) | 14.9 (2.4) | 0.2 (0.1) | 2.4 (1.6) | 1.4 (0.1) | 2.5 (0.7) | 4390 (741) | 6122 (414) | 0.8 (0.1) | 1.0 (0.2) | 226 (85) | 1923 (993) |
| **17** | 93.0 (0.9) | 82.2 (1.8) | 6.7 (0.9) | 14.9 (1.7) | 0.2 (0.1) | 2.9 (0.5) | 1.3 (0.1) | 2.6 (0.4) | 4561 (565) | 5944 (755) | 0.9 (0.2) | 1.2 (0.1) | 206 (61) | 2061 (284) |
| **18** | 93.3 (0.8) | 84.9 (0.4) | 6.5 (0.8) | 13.5 (0.5) | 0.2 (0.0) | 1.6 (0.2) | 1.4 (0.1) | 2.2 (0.2) | 4310 (432) | 6009 (395) | 0.8 (0.2) | 0.9 (0.2) | 164 (36) | 1513 (120) |

*Table 1: Activity metrics involving time spent above or below an acceleration threshold for DE50-MD and WT dogs at all ages. Results for metrics from left to right: % time spent at rest, % time spent at low intensity activity, average active bout duration at low intensity (s), total active bout count at low intensity activity, average bout duration at high intensity activity, total bout count at high intensity activity for DE50-MD and WT control dogs. Mean (standard deviation) displayed for each metric at each age between 3 and 18 months. N numbers varied at each age, see Fig. 1 for further details. Metrics counting discrete bouts have been rounded to the nearest integer, acceleration metrics ($MX_{ACC}$) to 2 decimal places. and all other metrics to 1 decimal place). Symbols indicate group effect at all ages unless highlighted: * p<0.05, ** p<0.01, *** p<0.001, $^{f}$ not significant at that age point.*

| Age in Months | Activity Intensity ($ms^{-1}$)*** | | $M2_{ACC}$ (g)** | | $M30_{ACC}$(g)** | | $M60_{ACC}$(g)** | | PC1*** | | PC2 | |
|---|---|---|---|---|---|---|---|---|---|---|---|---|
| | *DE50* | *WT* | *DE50* | *WT* | *DE50* | *WT* | *DE50* | *WT* | *DE50* | *WT* | *DE50* | *WT* |
| **3** | 7101 (894) | 9872 (1878) | 1.14 (0.11) | 1.63 (0.15) | 0.52 (0.02) | 0.72 (0.10) | 0.39 (0.02) | 0.53 (0.08) | -0.5 (1.0) | -3.6 (1.8) | 2.5 (0.5) | 2.0 (0.7) |
| **4** | 6151 (1082) | 10068 (1950) | 1.09 (0.12) | 1.7 (0.19) | 0.48 (0.10) | 0.79 (0.16) | 0.35 (0.08) | 0.56 (0.11) | 0.3 (1.2) | -3.9 (1.9) | 1.5 (0.6) | 1.2 (1.0) |
| **5** | 5088 (807) | 8759 (1718) | 0.96 (0.14) | 1.75 (0.36) | 0.38 (0.05) | 0.68 (0.14) | 0.29 (0.03) | 0.48 (0.10) | 1.4 (0.8) | -2.5 (1.8) | 1.2 (0.6) | 1.5 (0.7) |
| **6** | 4551 (382) | 8235 (1270) | 0.95 (0.08) | 1.65 (0.18) | 0.37 (0.04) | 0.67 (0.08) | 0.28 (0.03) | 0.47 (0.07) | 2.1 (0.5) | -2.1 (1.3) | 1.0 (0.3) | 1.2 (0.6) |
| **7** | 4251 (484) | 7303 (560) | 0.93 (0.15) | 1.63 (0.10) | 0.34 (0.03) | 0.63 (0.05) | 0.25 (0.02) | 0.44 (0.04) | 2.3 (0.6) | -1.3 (0.5) | 0.6 (0.3) | 0.9 (0.7) |
| **8** | 4225 (456) | 6591 (694) | 0.99 (0.14) | 1.53 (0.16) | 0.33 (0.05) | 0.57 (0.10) | 0.25 (0.03) | 0.39 (0.06) | 2.3 (0.8) | -0.5 (1.0) | 0.1 (0.5) | 0.7 (0.7) |
| **9** | 3823 (673) | 7320 (615) | 0.83 (0.14) | 1.66 (0.20) | 0.31 (0.05) | 0.63 (0.06) | 0.24 (0.04) | 0.43 (0.05) | 2.9 (0.8) | -1.2 (0.7) | 0.5 (0.4) | 0.7 (0.8) |
| **10** | 3825 (773) | 7735 (1325) | 0.91 (0.19) | 1.78 (0.34) | 0.33 (0.05) | 0.68 (0.15) | 0.24 (0.04) | 0.48 (0.11) | 2.7 (0.9) | -1.8 (1.8) | 0.1 (0.7) | 0.3 (1.3) |
| **11** | 3571 (322) | 7825 (1296) | 0.87 (0.13) | 1.74 (0.30) | 0.29 (0.03) | 0.67 (0.13) | 0.22 (0.02) | 0.46 (0.09) | 3.0 (0.4) | -1.9 (1.6) | -0.3 (0.6) | 0.5 (1.1) |
| **12** | 3358 (481) | 8262 (2124) | 0.86 (0.19) | 1.94 (0.57) | 0.28 (0.02) | 0.71 (0.19) | 0.21 (0.02) | 0.49 (0.14) | 3.1 (0.7) | -2.2 (2.4) | -0.5 (0.6) | 0.1 (1.3) |
| **13** | 3886 (607) | 7416 (1545) | 0.96 (0.20) | 1.77 (0.26) | 0.32 (0.05) | 0.66 (0.14) | 0.24 (0.03) | 0.44 (0.10) | 2.7 (0.8) | -1.5 (1.8) | 0.2 (0.4) | 0.0 (0.8) |
| **14** | 3257 (244) | 9229 (3026) | 0.86 (0.11) | 2.09 (0.32) | 0.29 (0.02) | 0.88 (0.24) | 0.21 (0.01) | 0.6 (0.20) | 3.3 (0.3) | -3.9 (3.5) | -0.3 (0.8) | -1.6 (1.3) |
| **15** | 3327 (455) | 7744 (926) | 0.83 (0.10) | 2.02 (0.28) | 0.29 (0.03) | 0.77 (0.08) | 0.21 (0.02) | 0.48 (0.06) | 3.3 (0.4) | -2.8 (1.0) | -0.2 (0.6) | -1.5 (1.3) |
| **16** | 3184 (410) | 8278 (2617) | 0.86 (0.13) | 2.16 (0.69) | 0.28 (0.02) | 0.76 (0.29) | 0.2 (0.02) | 0.53 (0.21) | 3.4 (0.5) | -2.4 (2.8) | -0.4 (0.6) | -0.9 (1.3) |
| **17** | 3220 (283) | 8671 (877) | 0.88 (0.11) | 2.14 (0.08) | 0.28 (0.02) | 0.9 (0.09) | 0.2 (0.01) | 0.57 (0.08) | 3.3 (0.3) | -3.5 (1.5) | -0.7 (0.7) | -1.7 (0.6) |
| **18** | 3081 (264) | 6948 (156) | 0.8 (0.06) | 1.9 (0.07) | 0.27 (0.02) | 0.66 (0.04) | 0.2 (0.02) | 0.42 (0.03) | 3.4 (0.3) | -1.3 (0.7) | -0.8 (0.9) | -0.7 (0.1) |

*Table 2: Activity intensity, $MX_{ACC}$, PC1 and PC2 for DE50-MD and WT dogs at all ages. Results for metrics from left to right: activity intensity, $M2_{ACC}$, $M30_{ACC}$, $M60_{ACC}$, PC1 and PC2 for DE50-MD and WT control dogs. Mean (standard deviation) displayed for each metric at each age between 3 and 18 months. N numbers varied at each age, see Fig. 1 for further details. Metrics counting discrete bouts have been rounded to the nearest integer, acceleration metrics ($MX_{ACC}$) to 2 decimal places and all other metrics to 1 decimal place). Symbols indicate group effect at all ages unless highlighted: * p<0.05, ** p<0.01, *** p<0.001,$^{f}$ not significant at that age point. PC1 and PC2 refer to principal components 1 and 2.*

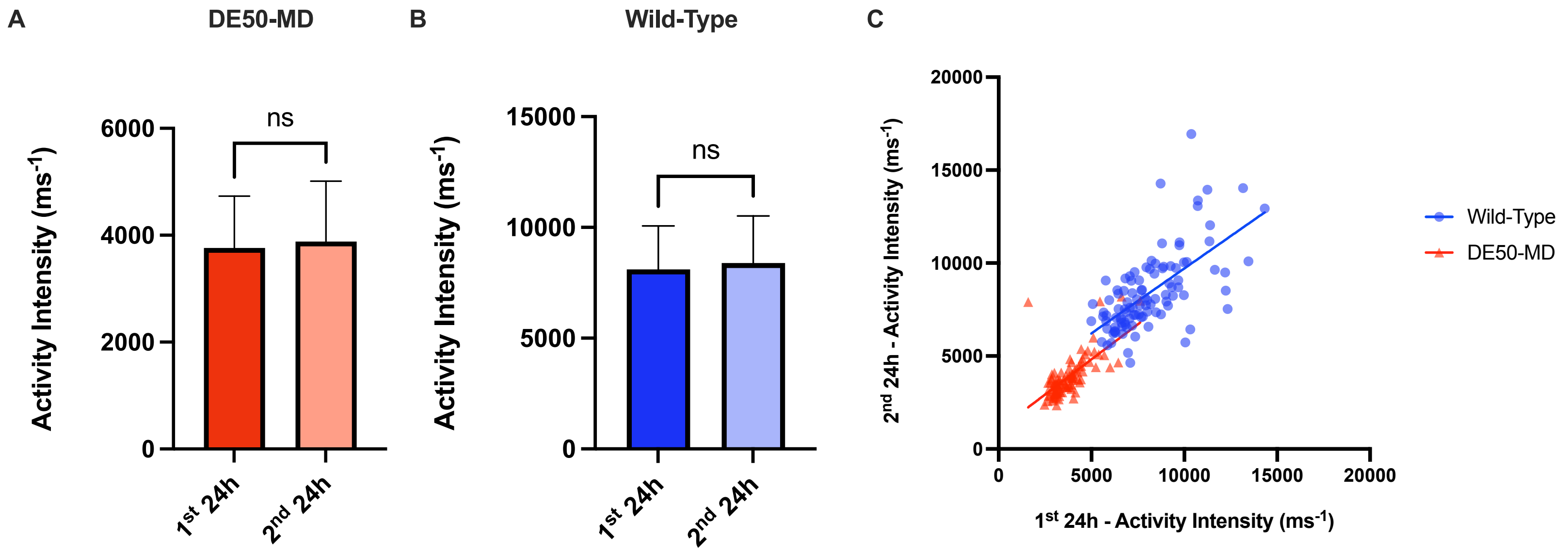


*Figure S1: Activity Intensity (mean and standard deviation) metrics computed for the 1st and 2nd 24h periods for **A**) Wild-Type dogs (blue; N=14 dogs; N=103 sets of 48h activity monitoring recordings in total; and **B**) DE50-MD dogs (red; N=11 dogs; N=95 sets of 48h activity monitoring recordings in total), and **C**) the correlation between the results for the two time periods. Note the lack of any significant difference in Activity Intensity (ns; P>0.05) in the second period compared with the first in either genotype. Recordings within each animal between first and second 24 hour period were highly correlated for each genotype (p<0.0001; WT: slope estimate: 0.75 +/- 0.09 SE; DE50-MD slope estimate: 0.69 +/- 0.08 SE).*
